# Age-Related Colonic Mucosal Microbiome Community Shifts in Monkeys

**DOI:** 10.1101/2020.05.17.100560

**Authors:** Ravichandra Vemuri, Chrissy Sherrill, Matthew Davis, Kylie Kavanagh

## Abstract

Recent evidence suggests that gut microbiome changes that occur with age impact the health of the host. While it is known that the gut microbiome and physiological systems interact, the relationship between the microbiome in an aged body system remains to be clearly defined, particularly in the context of inflammation. Therefore, we aimed to evaluate systemic inflammation and the mucosal microbiome in young and old female vervet monkeys. Ascending colon mucosal biopsies and blood samples from healthy young and old monkeys were collected for 16S rRNA gene sequencing and cytokine analyses, respectively. To demonstrate microbial co-occurrence patterns, we used Kendall’s tau correlation measure of interactions between microbes. We found elevated levels of plasma MCP-1 and CRP in old monkeys, which are indicative of higher systemic inflammation. Microbiome analysis revealed increases in abundance of opportunistic pathobionts such as members of the Proteobacteria phylum in old monkeys. At the family level, abundances of *Prevotellaceae* and *Helicobacteraceae* were higher in old monkeys than in young. We also found significantly lower Firmicutes to Bacteroidetes ratio (P= 0.03) and lower abundance of butyrate-producing microbes in old monkeys, consistent with a less healthy profile. Microbial community co-occurrence analysis revealed 13 nodes and 41 associations in the young monkeys, but only 12 nodes and 21 associations in the old monkeys. Our findings provided novel insights into systemic inflammation and gut microbial interactions, highlights the importance of the mucosal niche changes with age, and may facilitate further understanding of the decline in the stability of the microbial community with aging.

## Introduction

Aging is a complex, multifactorial process associated with the time-dependent decline of physiological functions, including gastrointestinal and immune systems. The interactions of these systems contributes to increased risk of age-related diseases (1, 2). Altered intestinal and immune function with aging, along with shifts in lifestyle and diet, can affect the composition and stability of the gut microbiome. The gut microbiome and its interactions within the host are suggested to be a vital factor in healthy aging. These interactions influence short-chain fatty acid production, maintenance of gut barrier integrity, regulation of immune responses, and provision of defense against pathogens (3).

Seminal studies on aging have revealed drastic changes in gut microbiome in early stages of life, adulthood, and old age (4-7). These studies provide evidence that the most dominant gut bacterial phyla are Bacteroidetes (Gram-negative), Firmicutes (Gram-positive) and Proteobacteria (Gram-negative). During aging, variable changes in phyla such as Bacteroidetes and Firmicutes, and an increase in pathobionts such as Proteobacteria are observed (4, 8). Although not consistent, such changes in the microbiome (dysbiosis) were linked to mucosal barrier dysfunction, inflammation, and disease susceptibility in animal studies (9-12). However, most studies on microbiota primarily characterize fecal microbiomes, as they can be sampled easily and non-invasively, but may not fully represent the mucosal microbiome, which directly interacts with intestinal epithelial cells and the immune system (13). A limited number of animal and human studies, predominantly focused on inflammatory bowel disease, along with a few on aging, have examined the mucosal-associated microbiome and found location-specific microbiome and inflammation in the intestine (14-18).

We previously demonstrated that reduced mucosal barrier function and inflammation leads to metabolic diseases in aged monkeys (12, 19, 20). Further, many studies show that fecal microbiomes do not change with aging or with diet, despite disparate health outcomes, which substantiates the need to evaluate microbial interactions closer to the sites of host-microbe cellular interactions (20-22).

The gut microbial composition is influenced by numerous factors, including diet, antibiotics, sex hormones, circadian rhythm, and age (3). Apart from these factors, ecological interactions between microbes also contribute essentially to microbial community changes (23-25). Microbes interact with each other (either compete or cooperate) to form an intricate biological network in the gut ecosystem (24). The nature of microbe-microbe interaction ranges from symbiotic to antagonistic, which determines the overall composition and function of the ecosystem. Although there are a few studies on microbial network analysis, there is insufficient exposition of the aging microbiome. Moreover, knowledge about the composition of microbial communities thus far is limited to taxonomic changes. Given the high variability and complex nature of the aging microbiome, not only taxonomic changes but also microbe-microbe interactions are essential to understand the dynamics of the gut microbial community.

To understand the complex nature of the aging microbiome and inflammation, we evaluated the non-human primate (NHP) colonic mucosal microbiome, due to the close resemblances of old world NHPs to human physiology (12, 20). The goal of the current study was to apply a novel ecological approach to understanding the colonic mucosal microbiome differences in aged NHPs known to have higher microbial translocation burdens (12), with the goal of providing novel insights into age-associated inflammation and gut microbial interactions to facilitate our understanding of the stability of the microbial community across age groups.

## Materials and Methods

### Animals

Study subjects (N=19) resided in a breeding colony of vervet monkeys (*Chlorocebus aethiops sabaeus*) at Wake Forest University School of Medicine. The demographic details of these subjects, and confirmation of their mucosal dysbiosis and intestinal leakiness, have been published (12). All procedures were conducted on a protocol approved by the Wake Forest University Institutional Animal Care and Use Committee according to guidelines from the Guide for Care and Use of Laboratory Animals (Institute for Laboratory Animal Research) and in compliance with the USDA Animal Welfare Act and Animal Welfare Regulations (Animal Welfare Act as Amended; Animal Welfare Regulations).

### Sample collection, DNA extraction, and 16s rRNA sequencing

Blood and tissue samples were collected at a predetermined necropsy endpoint. Samples collected for microbiome characterization included feces and a colonic mucosal scraping from the ascending colon. DNA was extracted from these samples (Qiagen, Hilden, Germany), and the quality of the DNA was determined via gel electrophoresis. Mucosally-sourced DNA was evaluated for estimated total bacterial gene content in duplicate via PCR and was normalized to the starting tissue weight. DNA was further quantitated and sequenced by a commercial laboratory (Hudson Alpha Institute for Biotechnology, Huntsville AL, USA) and taxonomic assignments were done as previously described (17).

### Circulating Biomarkers

Plasma was used to assess circulating monocyte chemoattractant protein (MCP)-1 and C-reactive protein (CRP) via ELISA following manufacturer’s instructions (R&D Systems, Minneapolis, MN, USA) (1).

### Microbiome data analysis

Bioinformatics analysis tools MicrobiomeAnalyst (26), MEGAN 6 (Community Edition) (27) and METAGENassist (28) were employed to quantitate the bacterial taxonomic profiles between groups. The taxonomic output reads (total: 441567) from all the samples were merged and converted into the widely accepted file formats .biom file and comma-separated values (CSV) file for downstream analyses. To obtain taxonomic profiles, this .biom file or .CSV file along with the metadata for each sample were imported, filtered, normalized and analyzed as previously described (29). Briefly, data were filtered by removing features with all zeros or containing very low counts to improve downstream analysis. After data filtering, data were normalized by scaling (unification of all samples to the same scale) and transformation (stabilization of variation) to avoid data sparsity issues and enable meaningful comparison. Alpha (α) diversity profiles to measure species richness and evenness were analyzed using Shannon’s and Simpson’s indexes. Further, a rarefaction curve analysis was performed to measure species richness alone. To determine the microbial variation (Beta (β) diversity) between the groups, we used clustering methods including partial least squares-discriminant analysis (PLS-DA) and principal component analysis (PCA). PLS-DA is a supervised classification method designed to enhance the separation between the groups by rotating the PCA components to achieve maximum separation. Because PLS-DA can overfit data and avoid false discovery regarding the microbial separation, we used 1000 permutations to validate these model as previously described (28, 29). PLS-DA and PCA associated bi-plots were generated to understand the impact on the division among groups as previously described. Additionally, principal coordinate analysis (PCoA) plots were generated with weighted and unweighted UniFrac β diversity metric.

### Functional metagenomics analysis

To obtain functional profiles of age-dependent gut microbiome changes, we used METAGENassist. METAGENassist uses automated taxonomic-to-phenotype mapping through a unique microbial phenotype database with information on more than 11,000 microbial species compiled from BacMap, NCBI and GOLD. The data was normalized by column-wise normalization (taxon vs taxon) and auto-scaling (mean-centered and divided by the standard deviation of each variable) by total abundance before calculating group means as previously described (30). For phenotype categories such as oxygen requirement and metabolism that may have multiple phenotypic traits associated with a given taxon, a bar chart was generated to represent the percentage of taxa with each trait. For each metabolic phenotype analysis, a supervised PLS-DA plot was used as described (28, 29).

We used Phylogenetic Investigation of Communities by Reconstruction of Unobserved States (PICRUSt) (30) and identified the Kyoto Encyclopedia of Genes and Genomes (KEGGs) module to perform advanced functional analysis. The .biom file containing reads with GreenGenes taxonomical classification was converted into a .CSV file containing KEGG orthologs, and clusters of ortholog genes (COG) annotated reads by assigning them to KEGG database using PICRUST. The stacked plot was made with data containing KO annotated read counts using MicrobiomeAnalyst for comparative analysis of functional profiles across an individual sample. Dendrogram analysis using Bray-Curtis index dissimilarity measure and Ward clustering algorithm was performed to identify the dissimilarity between samples across groups. The abundance was computed by the sum of all hits belonging to each category (KEGG/COG metabolic pathway) and normalized by the size of the category.

### Microbial co-occurrence network analysis

To understand the ecological interactions between microbes, we used MEGAN 6 Community Edition to make microbial co-occurrence network plots. Each network plot comprises of nodes and edges. Nodes represent the microbial taxa, and the size of the node (1 to 10) represents the strength of the association. Edges (interconnecting lines) represent the nature of association such as co-occurrence (green color; positive associations) or co-exclusion (red; negative associations). Microbial co-occurrences and co-exclusion plots were generated using Kendall’s tau coefficient of correlation method (more robust to outliers) as previously described (27, 31). This particular correlation method uses the following formula to calculate the associations: (C-D)/(C+D) where C is the number of concordant pairs, and D is the number of discordant pairs. The tau correlation coefficient returns a value ranging from -1 to 1, where all positive values represent co-occurrences and negative values represent co-exclusion. In this study, the overall network threshold with significance > 0.05, and the edge threshold of 70% was selected to control the false positive rate. The microbial network is measured and controlled by a number of parameters. The detection threshold percentage (at least 1%) sets a minimum count required for a taxon to be considered present in a sample. The minimum prevalence (5%) and max prevalence percentage (95%) parameters are used to set the minimum and maximum percentage of samples in which a taxon can occur, respectively, to have the taxon to be presented by a node in the network. The probability sets parameter defines the minimum probability that a co-occurrence between two taxa A and B must attain to be represented by an edge in the graph.

### Statistical analysis

All statistical analysis was performed using GraphPad Prism software version 8 (San Diego, CA, USA). Data from young and old animals were compared using Student’s unpaired T-tests. Statistical significance was set at α < 0.05. Where appropriate, reported p-values where corrected for multiple testing. Abundance values are reported as percentage abundances for the groups.

## Results

### Higher systemic inflammation in older monkeys

Demographic data of the animals evaluated in this study have been reported (12). Young monkeys (N = 10) were on average 11 years as compared to 22 years in the old (N = 9) group (p<0.001), but were weight matched (4.87 vs. 4.94 kg, p=0.45) (12). Elevated systemic inflammation, which we assessed via MCP-1 and CRP measurements, is considered a hallmark of aging. Old monkeys showed significantly higher plasma levels of both MCP-1 (p=0.02, **Figure 1A**) and CRP (p=0.03, **Figure 1B**).

**Figure 1.**
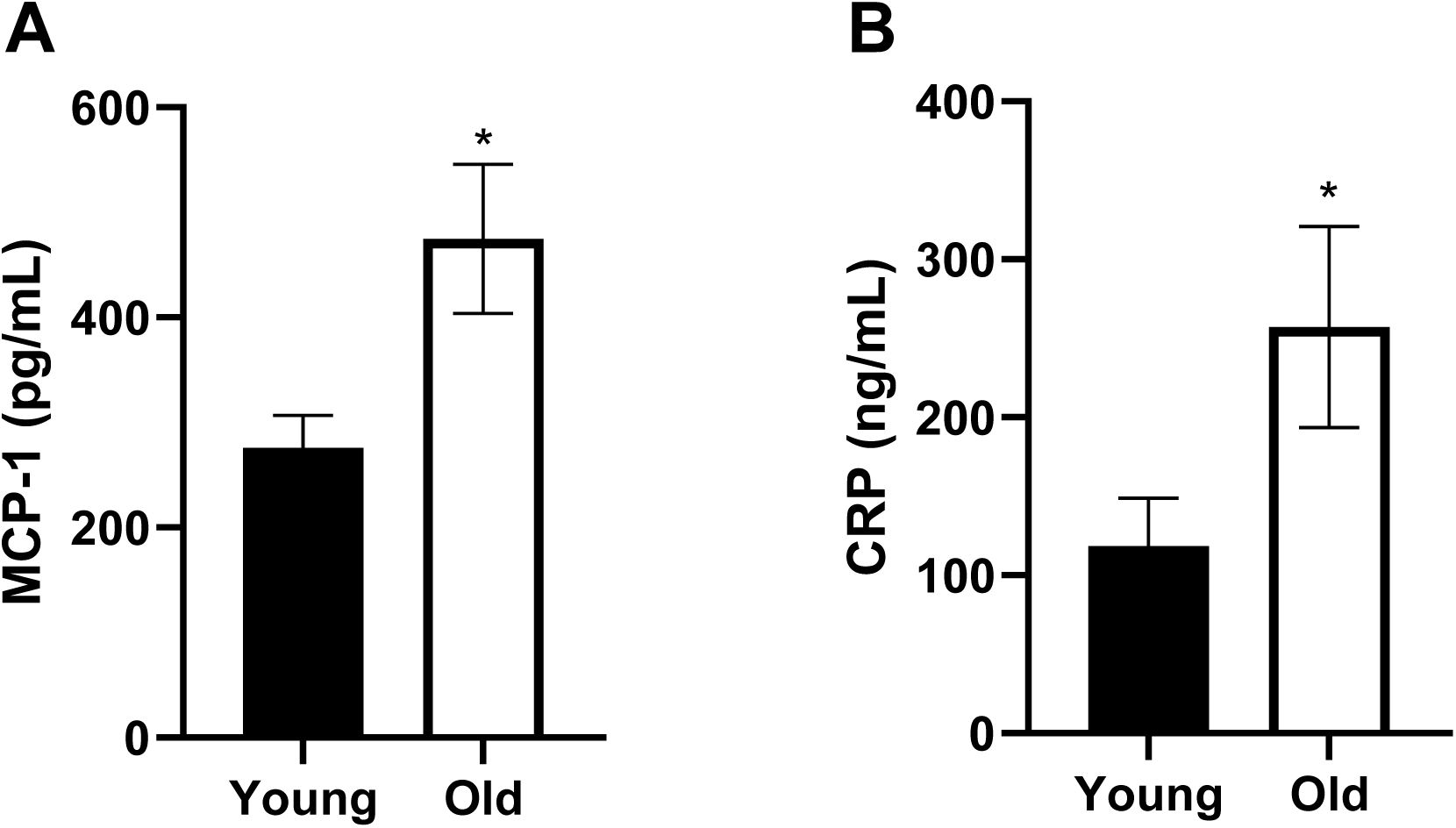
Higher systemic inflammation in old monkeys. Plasma levels of both MCP-1 (p=0.02) (**A**) and CRP (p=0.03) (**B**) were significantly higher in old monkeys when compared to young monkeys. The values are shown as the mean ± SEM. Significant differences with *p<0.05 were identified by *.

### Age-related changes in the colonic mucosal microbiome

We next analyzed microbiome profiles to investigate age-associated colonic mucosal changes in these animals. Beta (β) diversity profile using clustering methods, PLS-DA (**Figure 2A**) and PCoA (**Figure 2B**) of the unweighted UniFrac distance showed distinct separation in microbiome compositions between the young and old groups. PLS-DA plot divides two sets of samples with principal components 1 (percent variation: 13.8%), 2 (21%), and 3 (12.5%). Principal coordinates PC 1 (29.3%) PC 2 (24.1%) and PC3 (14.5%) of the PCoA plot separate most of the samples, though three outliers were observed in both groups. To evaluate the genera responsible for separation, we used 2D bi-plots based on beta diversity analysis (**Supplementary Figure 1A**). We observed no significant differences in alpha diversity by Shannon (**Supplementary Figure 1B**) and Simpson’s indexes (**Supplementary Figure 1C**), nor in rarefaction curve analysis for species richness and evenness (**Supplementary Figure 1D**). The most dominant bacterial phyla were Bacteroidetes, Firmicutes and Proteobacteria across the age spectrum (**Figure 2C and D**). However, the proportions of each taxon changed with age. In old monkeys, Bacteroidetes and Proteobacteria proportions increased by 9%, while Firmicutes decreased by 19% compared to young animals. Compared to the young group, the mucosal F/B ratio was significantly decreased in the old group (p=0.03) (**Figure 2E**), however it is notable that fecal F/B ratios were nearly identical between the two age groups (**Supplementary Figure 1E**) and illustrates the importance of sample type when evaluating age-related microbiome changes. At the family level, *Prevotellaceae, Ruminococcaceae* and *Helicobacteraceae* dominated across the age groups (**Figure 2F-G**). In old animals, proportions of both the families, *Prevotellaceae* and *Helicobacteraceae* increased when compared to young animals, though not achieving overall significance. Only two archaeal phyla were identified in our analysis, Euryarchaeota and Crenarchaeota, and both decreased with age (**Figure 2C-D**). In old monkeys, we also found reduced trends in genera *Butyricicoccus, Butyricimonas*, and *Butyrivibrio* though they did not reach overall significance (**Supplementary Figure 1F**).

**Figure 2.**
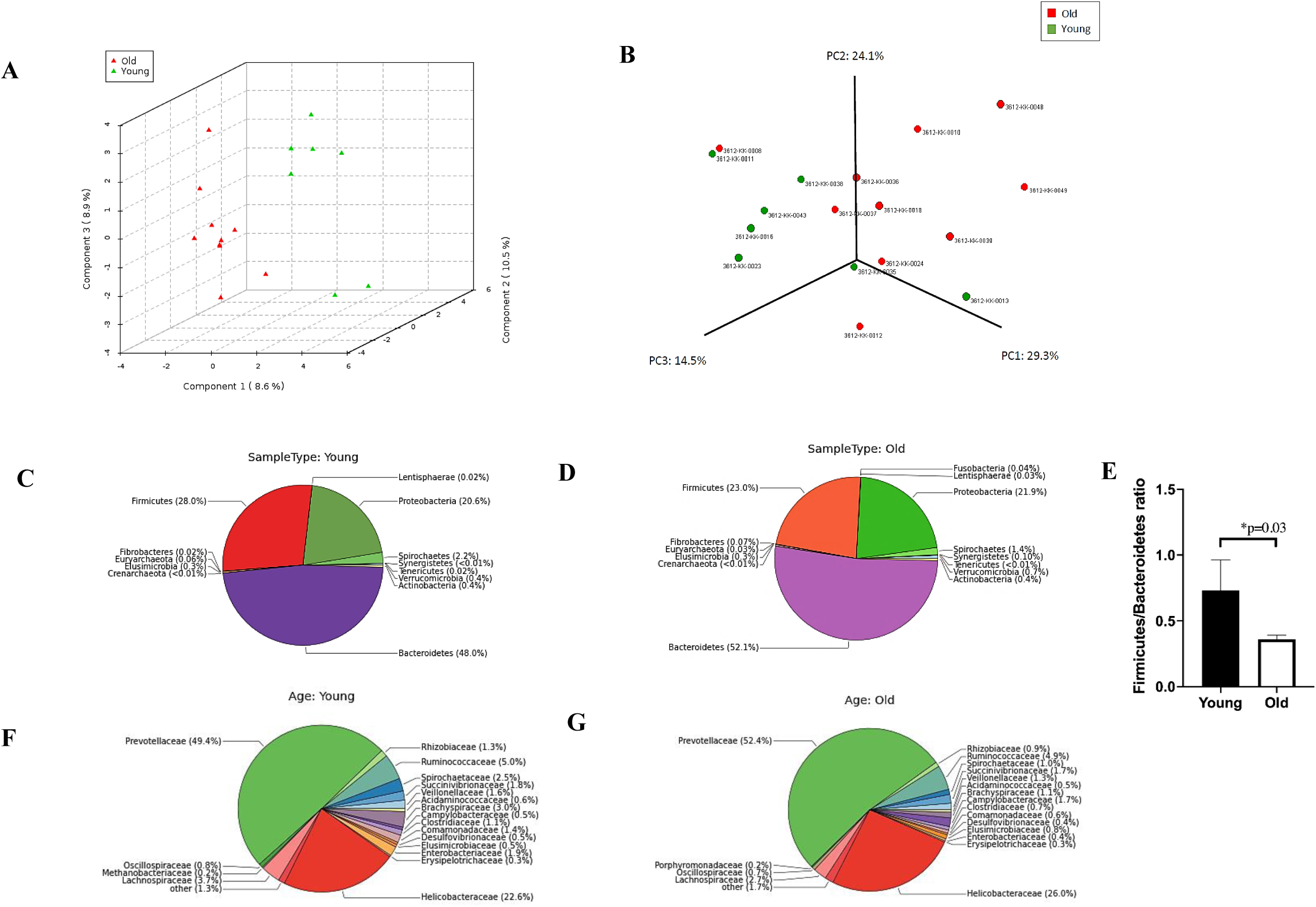
The gut microbiota analysis in young and old monkeys. (**A**) Partial least square discriminant analysis (PLS-DA) and (**B**) principal coordinate analysis (PCoA) (unweighted UniFrac) show divergence between the samples. The phylum level taxonomic composition in young (**C**) and old monkeys (**D**). (**E**) Firmicutes to Bacteroidetes ratio across age groups. The family level taxonomic compositions in young (**F**) and old monkeys (**G**). The values are shown as the mean ± SEM. Significant differences with * p< 0.05 were identified by *.

### Metagenomics functional analysis predicts metabolic changes with age

The taxonomic-to-phenotype mapping and visualization using METAGENassist revealed phenotypic changes in Gram staining and predicted metabolism. Gram phenotype showed a small increase in Gram-negative bacteria in old monkeys when compared to young monkeys (71.5% vs. 76.6%) (**Figure 3A and B**). With increased Gram-negative bacteria, we observed changes in microbial-associated metabolism, such as increased levels of ammonia oxidizers and nitrite reducers in the old group relative to the young group (**Figure 3C and D**). Further, PLS-DA cluster analysis on these metabolic phenotypic changes showed separation between groups (principal components 1 (29.1%), 2 (33.6%), and 3 (13.1%)), though outliers were observed (**Figure 3E**).

**Figure 3.**
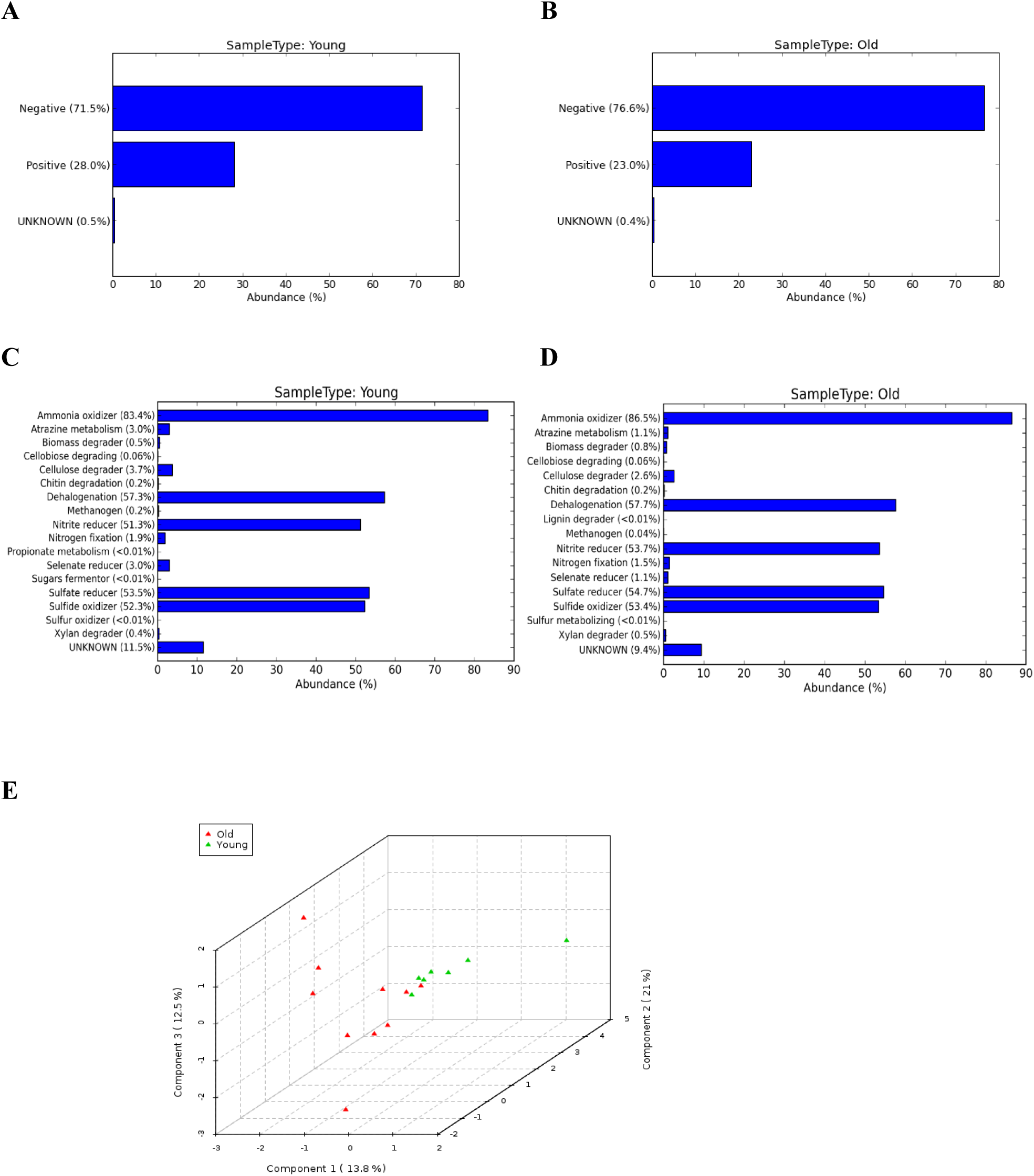
Metagenome function analysis between young and old monkeys. Gram-negative taxa showing differences in young (**A**) and old (**B**) monkeys, along with microbial metabolism changes between young (**C**) and old (**D**) monkeys. PLS-DA analysis (**E**) reveals distinct separation between the two groups.

KO annotated metagenomics sequences using MicrobiomeAnalyst led to the identification of KEGG- (**Supplementary Figure 2A**) and COG-predicted metabolic changes (**Supplementary Figure 2B**). At least 10 KO genes (predictive biomarkers) were identified as important by Random Forest analysis, which led to differences across the age groups (**Supplementary Figure 3A**). Although there were differences in actual abundances of a few pathways, such as carbohydrate metabolism (young: 13316.38 ± 21083.77, old: 20662.11 ± 10929.17), energy metabolism (young: 11218.90 ± 21089.17, old: 16039.93 ± 10215.94) and secondary metabolites production (young: 1617.14 ± 2831.19, old: 2635.45 ± 1857.97), none of these results attained significance (**Supplementary Table 1**). Dendrogram analysis by ward clustering showed a distinct separation between most of the samples (**Supplementary Figure 3B**), but again, there were no overall group delineations due to presence of three outliers.

### Microbial co-occurrence patterns in young and old monkeys

To understand the complex microbiome community structure and differences present in the study animals, we performed microbial co-occurrence analysis. This results in the formation of a biological network with taxonomic clusters (**Figure 4A and B**). This network is a combination of nodes as taxa and edges represent significant co-occurrence relationship between them. Network analysis revealed 13 nodes and 41 associations in the young group, and 12 nodes and only 21 associations in the old group (**Figure 4C**). Of 39 associations in the young group, 27 were co-occurrences, and 12 were co-exclusions. Conversely, we found only 14 co-occurrences and 7 co-exclusions in the old group, which can be visually appreciated as a simplified interactive landscape (**Figure 4B**). Summaries of these co-occurrences are shown in **Figure 4C** and **Supplementary Table 2**. Although Bacteroidetes was the most dominant phylum, it had only one association across age groups. The phyla with the highest numbers of co-occurrences in young animals were Euryarchaeota (Archaea), Synergistetes, Crenarchaeota (Archaea) and Actinobacteria, while Elusimicrobia phylum had most co-exclusions. Similarly, in old animals, Synergistetes had the most co-occurrences, and Elusimicrobia phylum had the most co-exclusions.

**Figure 4.**
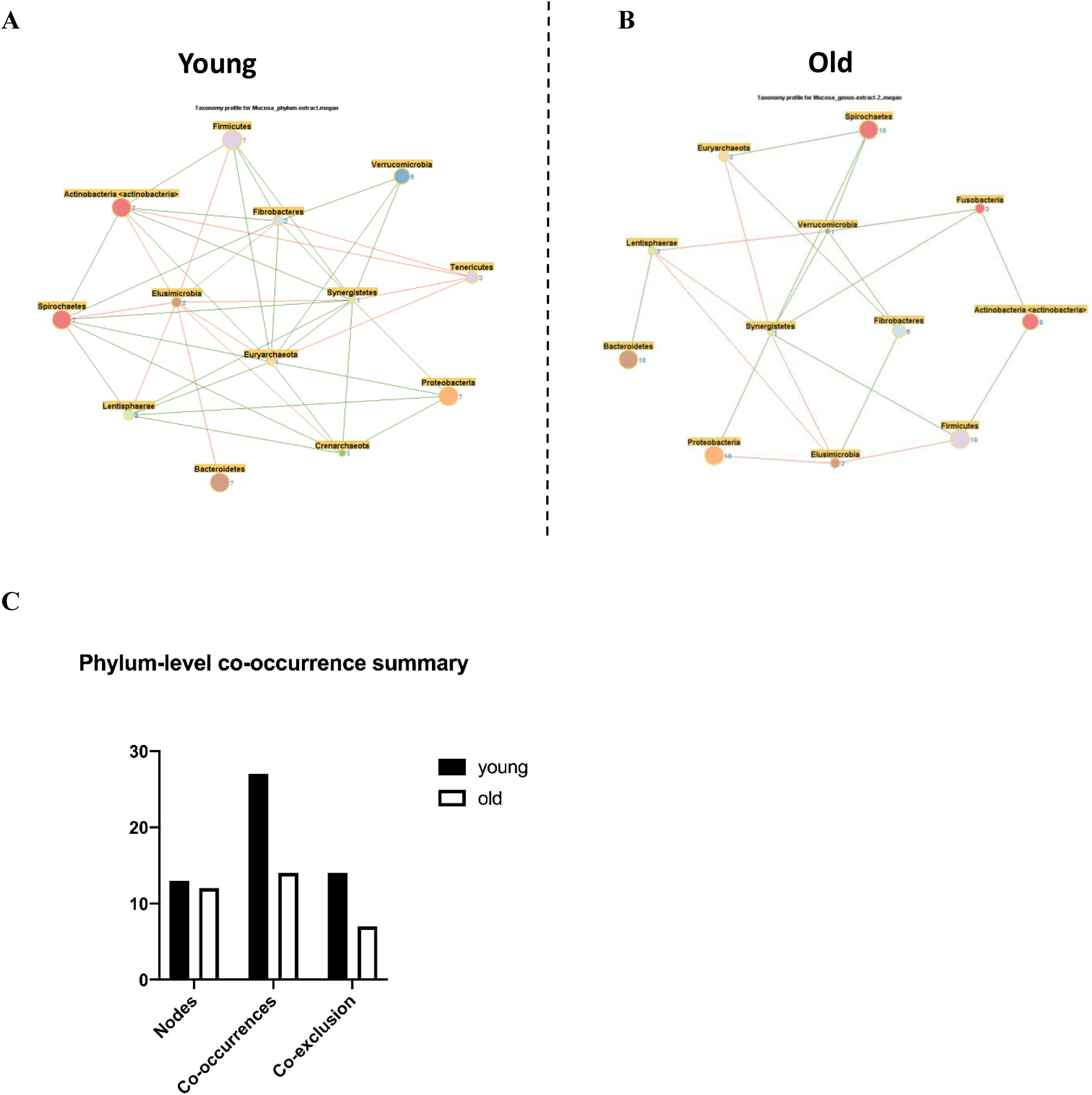
Microbial co-occurrence in the mucosal microbiome. Co-occurrence networks were constructed on the basis of the relative abundance profiles of colonic mucosal microbes using Kendall’s Tau correlation analysis in young (**A**) and in old (**B**) monkeys. Each node represents a phylum. Each edge indicates the sign of the association (green = positive [co-occurrences], red = negative [co-exclusions]). The thickness of the nodes represents the level of association between taxa. Summary (**C**) of co-occurrences and co-exclusion analysis.

## Discussion

In this study, we investigated the association between age-related changes in immune responses and gut microbiome, utilizing an NHP model. We found significant increases in plasma cytokine MCP-1 as well as CRP in older animals, together contributing to chronic systemic inflammation. Microbiome profiling revealed differences in gut microbial composition with age. Specifically, we observed increases in pathobionts phylum Proteobacteria and families *Prevotellaceae* and *Helicobacteriaceae* in old monkeys. We also found a significant reduction in Firmicutes/Bacteroidetes ratio with age and assessed microbe-microbe interactions utilizing novel microbial co-occurrence network analysis. The bacterial phyla, even with low abundances, had a significant impact on co-occurrence network and overall microbial communities.

The breach of mucosal epithelial integrity and leak of bacterial products in aging triggers aberrant inflammatory responses resulting in increased accumulation of pro-inflammatory mediators, thus leading to systemic inflammation (2, 10, 12, 20). Many systemic inflammatory cytokines and molecules, especially interleukin (IL)-6, MCP-1, IL-1β and CRP, have been considered as important factors associated with aging and age-related diseases (32-34). Once inflammation is triggered, pro-inflammatory IL-6 stimulates the production of CRP in the liver and subsequent release into the bloodstream (33). In the present study, we found significant increases in plasma cytokines in old animals which is consistent with previous studies of this animal cohort (12).

Multiple human and animals studies on aging have demonstrated alterations in gut microbiome, especially increases in pathogenic bacteria which are associated with inflammation (4, 5, 9, 10, 18). We found no difference in alpha (α) diversity among age groups, and these data were closely matched with those of a previous report (12). Our beta (β) diversity analysis shown in PCoA (unweighted UniFrac) and PLS-DA plots revealed distinct clustering of the samples. We observed noticeable changes in microbial taxonomic compositions between young and old animals. Consistent with studies on NHPs (35-37) and humans (4, 5, 7), we also found the gut microbiome was dominated by Bacteroidetes and Firmicutes phyla across the age groups. Similar to a study by Biagi et al. (4), we found an overall increase in opportunistic pathobionts such as Proteobacteria phylum, and changes in Bacteroidetes and Firmicutes in old monkeys. Decreased Firmicutes to Bacteroidetes (F/B) ratio with age has been linked to obesity and inflammation (38, 39). Confirming these findings, we observed a low F/B ratio in old monkeys and modulations in immune responses similar to humans.

Most of the common genera with high abundance in the old group were from families *Prevotellaceae* and *Helicobateraceae* (Proteobacteria). Human (7, 8, 40) and NHP (35, 37) studies reported prevalence of genera that belong to *Prevotellaceae* in the gut. Interestingly, members of *Prevotellaceae* and *Helicobacteraceae* are known to augment mucosal inflammation enabling dissemination of bacterial products to the systemic circulation (36, 37, 41). We previously demonstrated associations between increased systemic microbial translocation and innate immune responses impacting metabolic health in aged monkeys (12). This suggests a possible role of age-related microbiome changes in systemic inflammation and co-morbidities. In addition, butyrate, a microbial short chain fatty acid metabolite, is known to maintain immune homeostasis by instigating the release of anti-inflammatory IL-10 and activation of T regulatory cells (42, 43). Levels of butyrate and butyrate-producing bacteria are inversely proportional to age-related inflammation (43, 44). In line with these findings, we observed reduced trends in butyrate-producing bacterial genera such as *Butyricicoccus* spp., *Butyricimonas* spp., and *Butyrivibrio* spp. in the old monkeys, likely contributing to inflammation observed in this study (3, 45).

To gain functional insights into changes in the microbiome across age groups, we performed predictive metabolic profiling. This metabolic phenotypic analysis effectively adds to the existing scientific knowledge about known bacterial phenotypes. Gram-stain phenotypes appeared to determine the changes in microbial colonizing and associated metabolism. An increase in Gram-negative bacteria, such as Bacteroidetes and Proteobacteria phyla, were associated with increased ammonia oxidization and dehalogenation in old monkeys (37). Further, although not significant, many differences between young and old monkeys, such as carbohydrate metabolism, energy production, and secondary metabolites production, might be attributed to changes in the microbiome (46). Relative to differences observed in human populations, differences in metabolic phenotype were minimal in this study. A study by Rampelli et al. (6) reported more profound metabolic changes in centenarians than in older people (63-73 yrs.). The older animals in our study correspond to older people but not necessarily centenarian-like oldest old, which may explains the marginal metabolic changes observed in this study. However, the functional potential of gut microbiota clearly needs further validation.

We expanded our microbiome analysis beyond taxonomic composition changes by performing novel microbial co-occurrence analysis. New approaches to understanding the microbiome are required as the complexity of the network is poorly represented by individual abundances and ignores the niche of the host in which these abundances exist. Our major goal was to detect robust microbial associations and infer co-occurrence network patterns between microbes in a given environment within a relevant animal model whose microbiome has evolved under consistent external environmental conditions. In addition, understanding microbial interactions or interdependence in a gut microbial community will undoubtedly enhance our understanding of accurate microbial diversity (23, 24). Our co-occurrence network analysis revealed remarkable differences between young and old animals with a less-connected community observed in the mucosal niche of the aged colon. We found that the number of associations decreases with age, thus notably altering the microbiome. The dynamics of the microbiome and its overall structure were driven by the presence or absence of taxa and their associations. Bacteroidetes, being the most dominant and widely studied phylum, had the least associations. Conversely, less dominant phyla such as Euryarchaeota, Synergistetes, and Elusimicrobia had higher numbers of associations across both age groups. These findings suggest that the importance of absence or presence of particular genera, even with low abundance, can significantly impact the overall microbial community.

The major strength of this study is our characterization of colonic mucosal-associated microbiota across a wide spectrum of age in a translationally-relevant model. To our knowledge, this is the first study to evaluate such age-associated systemic inflammation and mucosal microbiome changes. The mucosa-associated microbiota provided novel information, especially compared with those of fecal microbiota, concerning host-microbe and microbe-microbe interactions. An important limitation of our report is the inclusion of only female animals. Although not consistently observed (35), studies in NHPs have previously shown sexual dimorphic microbiome and inflammation. We also acknowledge that our results can only inform regarding the colonic mucosal microbiota. The sample site was selected being the major microbial organ, which directly interacts with the immune system, and in the case of aging, we have consistently observed mucosal barrier deficits leading to microbial translocation biomarkers being elevated with aging in nonhuman primates (11, 12, 20, 22). Furthermore, co-occurrence network analysis provides important information on the complexity of microbial structure and their interaction patterns, but is limited in the ability to interpret the nature of these interactions. In spite of this limitation, microbial network analysis is an essential tool for theoretical proposition and hypothesis generation, which is a necessary precursor to experimental evidence. In this study, in order to generate the network plots, we employed a specific correlation method along with steps to eliminate any spurious findings. Although we utilized a single approach, it revealed remarkable patterns between community members and differences between young and old microbiomes. Overall, the approach enabled us to conclude that the gut microbes inclined to co-occur or co-exclude more than expected by chance.

## Conclusion

Although previous clinical studies have demonstrated changes in microbiome composition with age, many such results have been inherently confounded by variability in factors such as lifestyle, diet, and medications. Moreover, differences in fecal and mucosal microbiota sparked further inconsistency in results. Here, we utilized a NHP model that allowed us to mimic human conditions while also controlling the confounding factors. We observed age-related changes in inflammation and colonic microbiome between young and old animals. Increases in opportunistic pathogens(pro-inflammatory) belonging to Proteobacteria phylum and systemic inflammation with aging suggests the existence of comparable shifts between humans and NHPs. Further, we identified a peculiar microbial co-occurrence pattern with age, i.e., the taxa with low abundances had significant impact on altering the overall microbial structure. A more in-depth elucidation of the microbe-microbe interactions residing in the mucosa will significantly improve our understanding of the age-related changes in the microbiome and will guide the development of improved therapeutic strategies to maintain overall health in an aging population.

## Acknowledgements

We would like to thank the animal care, veterinary, and research staff at the Wake Forest University Primate Center as well as the NHPs themselves. RV performed data analysis and interpretation and participated in manuscript preparation. CS performed data analysis and interpretation and participated in manuscript preparation. KK designed the study, performed data analysis, supervised, and had overall oversight of the study and manuscript preparation.

