## Supplementary Figure 1 for "Age-Related Colonic Mucosal Microbiome Community Shifts in Monkeys"

**Supplementary Figure 1: The gut microbial diversity profiles of the colonic mucosal samples from young and old monkeys. A.** 2D Bi-plot showing the key taxa involved in PLS-DA based divergence**.** Alpha diversity plots using Shannon’s index **(B)**, Simpson’s index **(C)** and rarefaction **(D)** illustrate no difference in species evenness and richness. (**E**) Firmicutes to Bacteroidetes (Firm/Bact) ratio in fecal samples of young and old monkeys and (**F**) Decreased trends in butyrate-producing microbes in old monkeys.

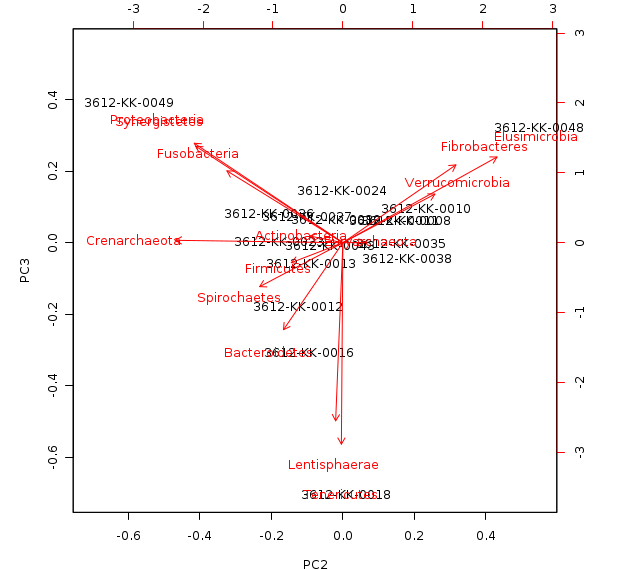
**A	B**	**C**

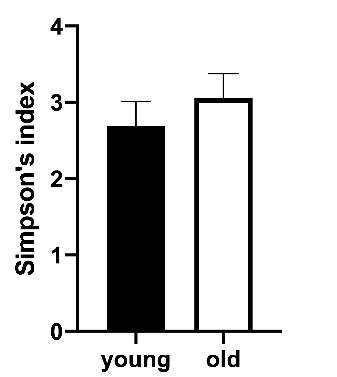

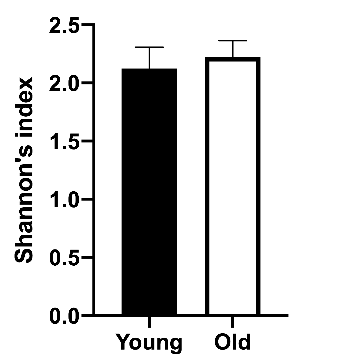

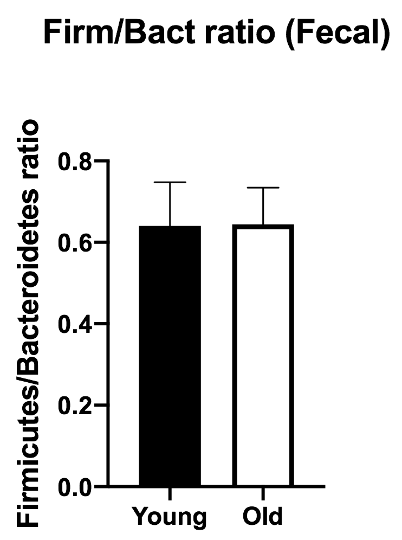
**D	E**

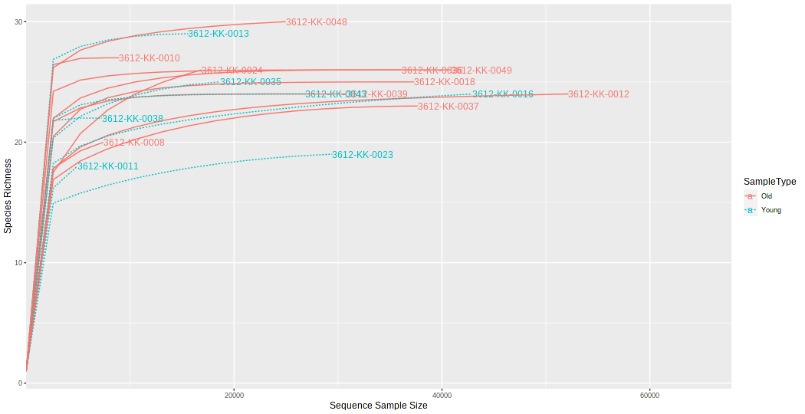

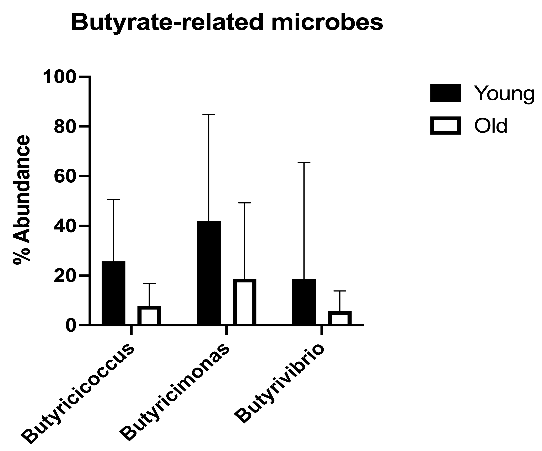
**F**
