## Supplementary Figure 2 for "Age-Related Colonic Mucosal Microbiome Community Shifts in Monkeys"

**Supplementary Figure 2: Metagenome function analysis differences between young and old monkeys.** Relative abundances of each metabolic pathway by KEGG metabolism (**A**) and COG metabolism (**B**) showed differences between groups in carbohydrate metabolism, energy metabolism and secondary metabolite production between young and old animals.

**A**

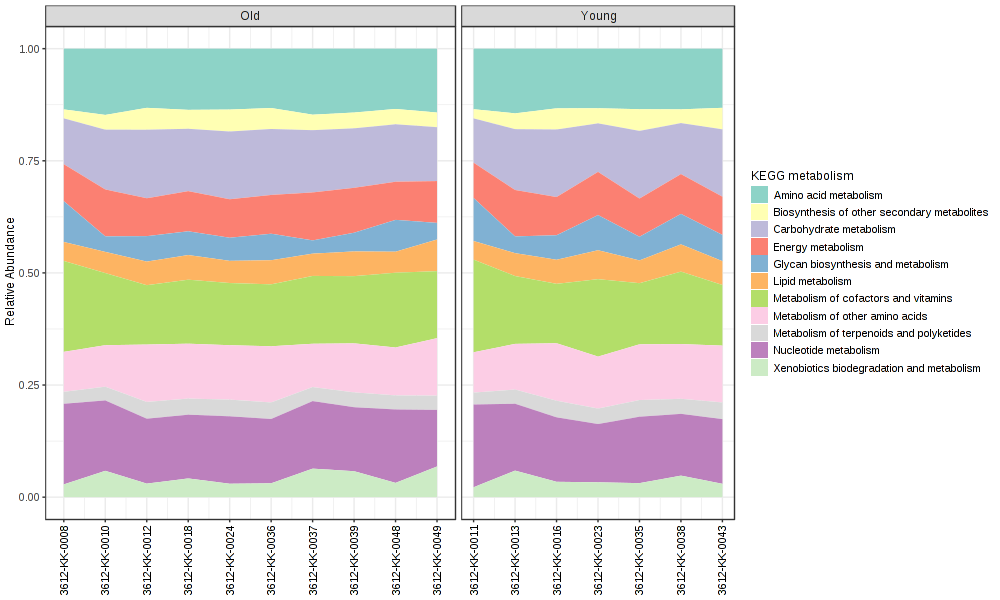

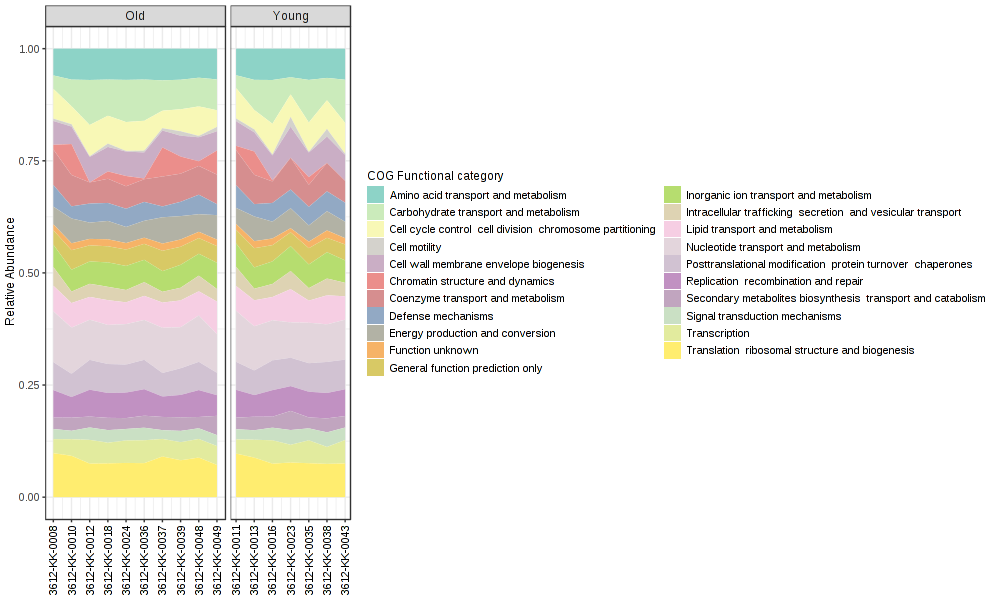
**B**
