## Supplementary Figure 3 for "Age-Related Colonic Mucosal Microbiome Community Shifts in Monkeys"

**Supplementary Figure 3**: Difference in predictive metabolic clustering between young and old monkeys. (**A)** Key compounds separating young and old based on variable importance in projection (VIP) score plot based on random forest analysis. (**B)** Dendrogram by Bray-Curtis dissimilarity and Ward clustering showing separation between samples.

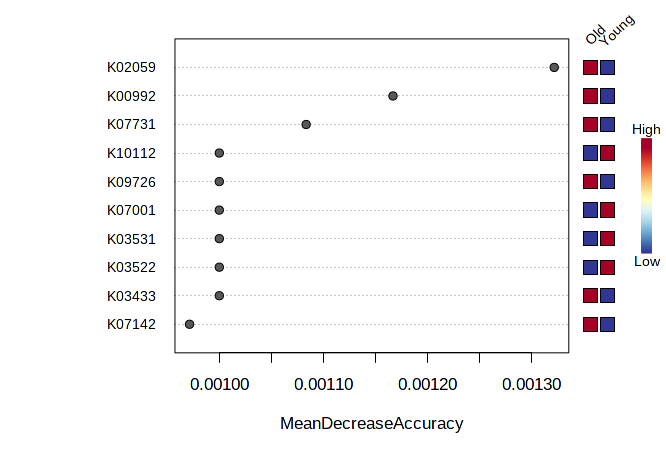
**A**

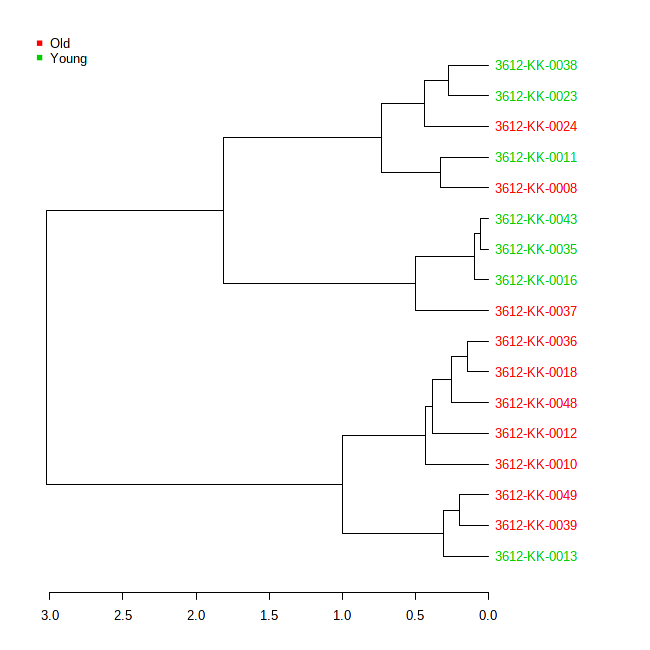

**B**
