## Supplementary Table 1 for "Age-Related Colonic Mucosal Microbiome Community Shifts in Monkeys"

**Supplementary Table 1**: List of predicted metabolic pathways based on KEGG metabolism in young and old monkeys

| **Age group** | Old | Old | Old | Old | Old | Old | Old | Old | Old | Old | Young | Young | Young | Young | Young | Young | Young |
| --- | --- | --- | --- | --- | --- | --- | --- | --- | --- | --- | --- | --- | --- | --- | --- | --- | --- |
| **Amino acid transport and metabolism** | 682 | 21524.5 | 14140 | 25824.5 | 2164 | 29488.5 | 8057 | 50994.5 | 27882.5 | 52653 | 1476 | 80122.5 | 6687.5 | 2722.5 | 7683 | 2635 | 7747.5 |
| **Carbohydrate transport and metabolism** | 287.8333 | 15481 | 16859.33 | 24909 | 2432.333 | 32435.33 | 6463.167 | 40099 | 23026 | 44628.17 | 593.8333 | 64272.5 | 7760.833 | 1295.333 | 8732.333 | 1643.833 | 8916 |
| **Cell cycle control cell division chromosome partitioning** | 138 | 2322 | 2552 | 4337 | 362 | 5176 | 832 | 6901 | 5139 | 5822 | 307 | 9359 | 1188 | 418 | 1307 | 499 | 1386 |
| **Cell motility** | 10 | 445.5 | 234.8333 | 1660.833 | 10.5 | 1477 | 208.8333 | 4019.833 | 245.5 | 3660.833 | 26.5 | 3202.833 | 122.5 | 799.5 | 88.33333 | 414.6667 | 259.5 |
| **Cell wall membrane envelope biogenesis** | 431 | 8255 | 7659.5 | 14072 | 1124.5 | 16547 | 2899.5 | 23913.5 | 15827.5 | 23913.5 | 971 | 33669 | 3567 | 1975.5 | 4030.5 | 1715 | 4300 |
| **Chromatin structure and dynamics** | 0.5 | 83 | 0 | 18 | 1.5 | 3 | 31.5 | 117.5 | 19 | 100 | 0.5 | 254.5 | 1 | 0 | 4 | 0 | 0 |
| **Coenzyme transport and metabolism** | 650.3333 | 15335.33 | 6610 | 14110 | 1082 | 14932.5 | 5331.5 | 32291.33 | 19528.67 | 36602.67 | 1401.5 | 52376.17 | 3178.167 | 2033.5 | 3767 | 1808.5 | 3755 |
| **Defense mechanisms** | 75 | 1163 | 1155 | 2041 | 170 | 2434 | 380 | 3243 | 2528 | 2427 | 170 | 4378 | 537 | 252 | 615 | 245 | 640 |
| **Energy production and conversion** | 453.3333 | 17213.33 | 7047.5 | 14908.5 | 1078.5 | 15591.5 | 6659 | 37783.33 | 16655.17 | 43009.17 | 905.5 | 62797.17 | 3452.667 | 1883 | 3851 | 1703.5 | 3939.5 |
| **Function unknown** | 302 | 9887 | 5834 | 12233 | 899 | 11890 | 3757 | 25289 | 12029 | 22509 | 641 | 37351 | 2869 | 928 | 3224 | 1477 | 3146 |
| **General function prediction only** | 739.1667 | 27745.83 | 14414.33 | 27313.5 | 2261.833 | 30449.33 | 10574.67 | 60760.33 | 30063.67 | 57661.33 | 1521.333 | 100869.7 | 6890 | 2794.833 | 7966.333 | 2703.833 | 7919.5 |
| **Inorganic ion transport and metabolism** | 420 | 10712 | 7367.833 | 14964.83 | 1216.5 | 15496.5 | 3663.833 | 27149.83 | 15382.5 | 31263.33 | 965 | 38564.83 | 3528 | 1663.167 | 4255.333 | 1772.667 | 4057.5 |
| **Intracellular trafficking secretion and vesicular transport** | 188 | 3290 | 2474.833 | 4824.833 | 352.5 | 5642 | 1148.833 | 9129.833 | 6003.5 | 9185.833 | 419.5 | 12512.33 | 1162 | 842.5 | 1281.833 | 668.1667 | 1423.5 |
| **Lipid transport and metabolism** | 215.5 | 5773 | 3150 | 6588 | 464 | 7111 | 2167.5 | 14568.5 | 7440.5 | 18636 | 452.5 | 22290.5 | 1537.5 | 1001.5 | 1675 | 854.5 | 1789 |
| **Nucleotide transport and metabolism** | 461.8333 | 11528.5 | 6357.333 | 11378.5 | 989.3333 | 13229.33 | 4118.667 | 23669.5 | 15792.5 | 24111.67 | 1005.333 | 40379 | 2998.833 | 1117.833 | 3510.333 | 1156.833 | 3480 |
| **Posttranslational modification protein turnover chaperones** | 414 | 9441 | 7548.167 | 13724.17 | 1095 | 15749.67 | 3493.667 | 25234.5 | 15518.67 | 22638.17 | 899 | 36727.17 | 3575 | 1517.833 | 3960.833 | 1594.833 | 4136.333 |
| **Replication recombination and repair** | 558 | 11512.5 | 9419 | 16227.5 | 1388 | 19538 | 4134.5 | 28816.5 | 20332.5 | 29132.5 | 1231 | 43942.5 | 4406 | 1772.5 | 4951.5 | 1775 | 5169.5 |
| **RNA processing and modification** | 0 | 6 | 1 | 43 | 0 | 0 | 4 | 97 | 0 | 88 | 0 | 70 | 3 | 1 | 1 | 11 | 0 |
| **Secondary metabolites biosynthesis transport and catabolism** | 75.5 | 2262 | 1147.5 | 2467 | 177 | 2717 | 828.5 | 5501 | 2681.5 | 8497.5 | 166.5 | 8539 | 549 | 448 | 629 | 315.5 | 673 |
| **Signal transduction mechanisms** | 115.5 | 2976.5 | 2765.5 | 5481.5 | 390 | 6096.5 | 1152 | 10023.5 | 4847.5 | 10595 | 253.5 | 12838 | 1344.5 | 818 | 1460 | 717.5 | 1558 |
| **Transcription** | 196.5 | 6766.5 | 6206.333 | 10045.33 | 904 | 12579.83 | 2650.833 | 17461.5 | 10176.83 | 19766.83 | 431 | 27081.83 | 2897.5 | 937.5 | 3272.667 | 881.1667 | 3364.167 |
| **Translation ribosomal structure and biogenesis** | 1058 | 27397.5 | 14120 | 26226 | 2224.5 | 30071 | 9833.5 | 56594 | 35771 | 54848.5 | 2278.5 | 95998.5 | 6674 | 2945 | 7829 | 2745.5 | 7813 |
