## Supplementary Table 2 for "Age-Related Colonic Mucosal Microbiome Community Shifts in Monkeys"

**Supplementary Table 2:** Summary of co-occurrences between young and old monkeys based on Kendall's Tau correlation analysis

| **Young** |  |  |  |
| --- | --- | --- | --- |
| Taxon1 | Taxon2 | Correlation (p<0.05) | Association |
| Bacteroidetes | Elusimicrobia | -1 | Negative |
| Actinobacteria | Firmicutes | 0.6 | Positive |
| Actinobacteria | Spirochaetes | 0.6 | Positive |
| Actinobacteria | Fibrobacteres | 1 | Positive |
| Actinobacteria | Elusimicrobia | 1 | Positive |
| Actinobacteria | Tenericutes | -0.7 | Negative |
| Actinobacteria | Fibrobacteres | 1 | Positive |
| Actinobacteria | Euryarchaeota | 1 | Positive |
| Actinobacteria | Synergistetes | 1 | Positive |
| Firmicutes | Actinobacteria | 0.6 | Positive |
| Firmicutes | Elusimicrobia | -0.6 | Negative |
| Firmicutes | Euryarchaeota | 0.5 | Positive |
| Firmicutes | Fibrobacteres | 0.8 | Positive |
| Firmicutes | Synergistetes | 0.7 | Positive |
| Elusimicrobia | Actinobacteria | -0.5 | Negative |
| Elusimicrobia | Firmicutes | -1 | Negative |
| Elusimicrobia | Fibrobacteres | -0.8 | Negative |
| Elusimicrobia | Euryarchaeota | -0.4 | Negative |
| Elusimicrobia | Crenarchaeota | -0.6 | Negative |
| Elusimicrobia | Bacteroidetes | -0.6 | Negative |
| Elusimicrobia | Lentisphaerae | -1 | Negative |
| Elusimicrobia | Spirochaetes | -1 | Negative |
| Proteobacteria | Synergistetes | 0.6 | Positive |
| Proteobacteria | Euryarchaeota | 0.7 | Positive |
| Proteobacteria | Lentisphaerae | 0.6 | Positive |
| Proteobacteria | Crenarchaeota | 1 | Positive |
| Euryarchaeota | Firmicutes | 0.5 | Positive |
| Euryarchaeota | Fibrobacteres | 1 | Positive |
| Euryarchaeota | Verrucomicrobia | 1 | Positive |
| Euryarchaeota | Synergistetes | 1 | Positive |
| Euryarchaeota | Lentisphaerae | 1 | Positive |
| Euryarchaeota | Elusimicrobia | 1 | Positive |
| Euryarchaeota | Proteobacteria | 0.6 | Positive |
| Euryarchaeota | Crenarchaeota | 1 | Positive |
| Euryarchaeota | Tenericutes | -1 | Negative |
| Fibrobacteres | Firmicutes | 0.8 | Positive |
| Fibrobacteres | Synergistetes | 1 | Positive |
| Fibrobacteres | Verrucomicrobia | 1 | Positive |
| Fibrobacteres | Euryarchaeota | 1 | Positive |
| Fibrobacteres | Spirochaetes | 0.6 | Positive |
| Fibrobacteres | Tenericutes | -1 | Negative |
| Fibrobacteres | Elusimicrobia | -1 | Negative |
| Verrucomicrobia | Fibrobacteres | 1 | Positive |
| Verrucomicrobia | Euryarchaeota | 1 | Positive |
| Verrucomicrobia | Synergistetes | 1 | Positive |
| Verrucomicrobia | Actinobacteria | 0.5 | Positive |
| Tenericutes | Fibrobacteres | -0.7 | Negative |
| Tenericutes | Actinobacteria | -1 | Negative |
| Tenericutes | Synergistetes | -1 | Negative |
| Tenericutes | Euryarchaeota | -1 | Negative |
| Spirochaetes | Actinobacteria | 0.6 | Positive |
| Spirochaetes | Fibrobacteres | 0.8 | Positive |
| Spirochaetes | Euryarchaeota | 1 | Positive |
| Spirochaetes | Synergistetes | 1 | Positive |
| Spirochaetes | Crenarchaeota | 0.6 | Positive |
| Spirochaetes | Lentisphaerae | 0.8 | Positive |
| Spirochaetes | Elusimicrobia | -1 | Negative |
| Lentisphaerae | Spirochaetes | 0.8 | Positive |
| Lentisphaerae | Synergistetes | 1 | Positive |
| Lentisphaerae | Euryarchaeota | 0.6 | Positive |
| Lentisphaerae | Proteobacteria | 0.6 | Positive |
| Lentisphaerae | Crenarchaeota | 0.8 | Positive |
| Lentisphaerae | Fibrobacteres | 0.5 | Positive |
| Lentisphaerae | Elusimicrobia | -1 | Negative |
| Synergistetes | Verrucomicrobia | 1 | Positive |
| Synergistetes | Firmicutes | 0.5 | Positive |
| Synergistetes | Fibrobacteres | 1 | Positive |
| Synergistetes | Euryarchaeota | 1 | Positive |
| Synergistetes | Spirochaetes | 1 | Positive |
| Synergistetes | Elusimicrobia | -1 | Negative |
| Synergistetes | Tenericutes | -1 | Negative |
| Synergistetes | Proteobacteria | 0.6 | Positive |
| Synergistetes | Crenarchaeota | 1 | Positive |
| Synergistetes | Actinobacteria | 1 | Positive |
| Synergistetes | Lentisphaerae | 1 | Positive |
| Crenarchaeota | Lentisphaerae | 0.8 | Positive |
| Crenarchaeota | Spirochaetes | 0.6 | Positive |
| Crenarchaeota | Elusimicrobia | -0.6 | Negative |
| Crenarchaeota | Euryarchaeota | 1 | Positive |
| Crenarchaeota | Synergistetes | 1 | Positive |
| Crenarchaeota | Proteobacteria | 1 | Positive |
| **Old** |  |  |  |
| Taxon1 | Taxon2 | Correlation (p<0.05) | Association |
| Bacteroidetes | Lentisphaerae | 0.7 | Positive |
| Firmicutes | Elusimicrobia | -0.6 | Negative |
| Firmicutes | Synergistetes | 0.8 | Positive |
| Firmicutes | Actinobacteria | 0.7 | Positive |
| Actinobacteria | Fusobacteria | 1 | Positive |
| Actinobacteria | Firmicutes | 0.7 | Positive |
| Proteobacteria | Synergistetes | 0.6 | Positive |
| Proteobacteria | Elusimicrobia | -1 | Negative |
| Synergistetes | Proteobacteria | 0.6 | Positive |
| Synergistetes | Spirochaetes | 1 | Positive |
| Synergistetes | Verrucomicrobia | 0.6 | Positive |
| Synergistetes | Fusobacteria | 1 | Positive |
| Synergistetes | Firmicutes | 0.8 | Positive |
| Synergistetes | Elusimicrobia | -0.6 | Negative |
| Synergistetes | Lentisphaerae | -1 | Negative |
| Synergistetes | Euryarchaeota | -1 | Negative |
| Fusobacteria | Verrucomicrobia | 0.7 | Positive |
| Fusobacteria | Synergistetes | 1 | Positive |
| Fusobacteria | Actinobacteria | 1 | Positive |
| Verrucomicrobia | Spirochaetes | 0.8 | Positive |
| Verrucomicrobia | Fibrobacteres | 1 | Positive |
| Verrucomicrobia | Synergistetes | 0.6 | Positive |
| Verrucomicrobia | Fusobacteria | 0.7 | Positive |
| Verrucomicrobia | Lentisphaerae | -1 | Negative |
| Spirochaetes | Euryarchaeota | 0.6 | Positive |
| Spirochaetes | Verrucomicrobia | 0.8 | Positive |
| Spirochaetes | Synergistetes | 1 | Positive |
| Euryarchaeota | Spirochaetes | 0.6 | Positive |
| Euryarchaeota | Fibrobacteres | 0.6 | Positive |
| Euryarchaeota | Synergistetes | -1 | Negative |
| Fibrobacteria | Verrucomicrobia | 1 | Positive |
| Fibrobacteria | Euryarchaeota | 0.6 | Positive |
| Fibrobacteria | Elusimicrobia | 0.8 | Positive |
| Lentispaerae | Bacteroidetes | 0.7 | Positive |
| Lentispaerae | Elusimicrobia | -1 | Negative |
| Lentispaerae | Synergistetes | -1 | Negative |
| Lentispaerae | Verrucomicrobia | -1 | Negative |
| Elusimicrobia | Proteobacteria | -1 | Negative |
| Elusimicrobia | Lentisphaerae | -1 | Negative |
| Elusimicrobia | Synergistetes | -0.6 | Negative |
| Elusimicrobia | Firmicutes | -0.6 | Negative |
| Elusimicrobia | Fibrobacteres | 0.8 | Negative |
